# The inverse relationship between fibronectin and neuroligins in the embryonic rat colon

**DOI:** 10.1101/759266

**Authors:** Ni Gao, Peimin Hou, Qiangye Zhang, Weijing Mu, Jian Wang, Dongming Wang, Aiwu Li

## Abstract

Our previous study identified that the abnormal expression of fibronectin (FN), neuroligin-1 (NL1) and neuroligin-2 (NL2) in the colons of children with Hirschsprung disease (HSCR), but the correlated relationship between the three in the development of the enteric nervous system (ENS) remains unclear. Colons of Wistar rats and PC12 neurons were used to investigate the relationship between FN and the neuroligins (NLs). Colon tissues from thirty healthy embryonic rats, including fifteen at embryonic day 16 (E16), eight at E18, seven at E20, and fifteen newborn rats within 24 hours (Ep0) were analyzed to determine the correlated expression of FN and NLs using Western blot (WB) analysis and real-time fluorescence quantitative PCR (qRT-PCR) methods. Small interfering ribonucleic acid (siRNA) targeting and gene plasmids were used to explore the functional interaction between FN and NLs by using PC12 neuron cells. Furthermore, we used recombinant FN and NL proteins to confirm their interactions. Our studies showed that there were downregulatory effects between FN and the NLs in the embryonic rat colon and different cell lines, indicating that FN and NLs could directly regulate each other, and there is a negative linear correlation between them. The imbalanced interaction between extracellular matrix and synapse-related genes may provide a new perspective for the pathogenesis and treatment of HSCR and neuronal intestinal malformations (NIMs).

## 1. Introduction

Neuroligins (NLs), which are expressed predominantly at the postsynaptic terminal, interact with neurexins (NXs) across the synaptic cleft and play critical roles in proper synapse maturation, function and neuronal function.[1] Limited research has shown that NLs may be expressed in the enteric nervous system (ENS). The extracellular matrix (ECM) is secreted by cells and composed of proteins and glycosaminoglycans (GAGs).[2] ECM constitutes the cell microenvironment and plays critical roles in guiding neighboring cell behaviors, such as migration, shape, survival, differentiation, and proliferation.[3–9] Fibronectin (FN) is one of the most vital ECM glycoproteins produced by mesenchymal cells in the colon and plays a crucial role in modulating the neural crest cell response.[10, 11] ECM homeostasis imbalance might affect the microenvironment surrounding enteric neural crest cells (eNCCs), which may be responsible for innervation deficiency due to incomplete colonization of eNCCs in the gut in patients with Hirschsprung disease.[12] Hirschsprung disease (HSCR) is attributable to failed migration of neural crest cells in the distal bowel and extending proximally for varying distances from the anus. HSCR causes a life-threatening bowel obstruction in approximately 1 in 5000 live births.^1^ The earlier the arrest of the migration is, the longer the aganglionic segment. We have found a temporal trend in the abnormal expression of FN, neuroligin-1 (NL1) and neuroligin-2 (NL2) in the diseased intestine of children with HSCR.[13, 14] These pieces of evidence led us to hypothesize that the abnormal development of FN, NL1 and NL2 may lead to intestinal lesions but does not explicitly indicate whether there is a real interaction between FN and NLs.

Therefore, we further investigated the expression of FN, NL1 and NL2 on different embryonic days and the relationship between them and confirmed the interaction between FN and NLs in this study. This work may contribute to the understanding of mechanistic research and confirms that synapse-related genes interact with the ECM microenvironment.

## Materials and methods

### Animals

Thirty healthy Wistar rats at embryonic days 16 (E16), 18 (E18), 20 (E20) and fifteen newborns within 24 hours (Ep0) were used in this study. All Wistar rats were purchased from the animal facility of Shandong University. All procedures were approved by the local ethics committee and were performed at Qilu Hospital, Shandong University. Rats were anesthetized with chloral hydrate and then sacrificed by cervical dislocation. Segments of the full thickness colon, approximately 1.0 cm distal to the ileocecal junction, were harvested. The entire surgery was performed on ice, and the colon specimens were stored at −80°C.

### Cell culture, transfection, and treatments

293T cells (a human embryonic renal epithelial cell line that is widely used in cell biology for their reliable growth and propensity for transfection)[15] and the PC12 cell line (a pheochromocytoma of the rat adrenal medulla derived from neural crest cells, which was widely used in studies of neuronal disease and in *vitro* neurobiological studies),[16] obtained from the cell bank of the Institute of Biochemistry and Cell Biology (Shanghai, China), were cultured in Dulbecco’s Modified Eagle’s Medium (DMEM, Gibco) supplemented with 10% fetal bovine serum (FBS, Gibco). Cells were then kept in a humidified incubator at 37°C in a 5% CO_2_ atmosphere for either 24 or 48 hours to reach 80% confluence in six-well plates. After transfection or stimulation with RGD (0, 5, 10 and 20 μg/mL) and NL recombinant proteins (NLsrp) (0, 0.2, 1 and 5 μM) for 30 min, cells were harvested for biochemical analyses.

Transfections were carried out with Lipofectamine 2000 (Invitrogen) as described in the manufacturer’s protocols.

### Sequences of siRNA vectors

FN1 (Rattus norvegicus): GCUAUUACAGAAUCACCUATT and UAGGUGAUUCUGUAAUAGCTT.

NL1 (Rattus norvegicus): CCAGCUGGGCUGUUAGUUUTT and AAACUAACAGCCCAGCUGGTT.

NL2 (Rattus norvegicus): GCUGUUCCAGAAGGCCAUUTT and AAUGGCCUUCUGGAACAGATT.

Negative control: UUCUCCGAACGUGUCACGUTT and ACGUGACACGUUCGGAGAATT.

### Antibodies and reagents

Detailed information on the antibodies and primers is listed in Tables 1 and 2. The other reagents for Western blot (WB) included Protein Extraction Kit (Beyotime, China), BCA Protein Concentration Determination Kit (Beyotime, China) and SDS-PAGE Gel Preparation Kit (Beyotime, China). Reagents for qRT-PCR included TRIzol (Ambion, USA), PrimeScript™ RT Master Mix (Perfect Real Time) (TaKaRa, Japan), and SYBR Premix Ex Taq™ Tli RNaseH Plus (TaKaRa, Japan).

**TABLE 1:** Detailed information of antibodies.

| Antigen | Antibody | Dilution | Application | Source |
| --- | --- | --- | --- | --- |
| Neurologin-1 | Mouse monoclonal | 1/1000 | Detect NL1 with Western blot | Abcam, USA |
| Neurologin-2 | Goat polyclonal | 1/1000 | Detect NL2 with Western blot | Abcam, USA |
| Fibronectin | Rabbit polyclonal | 1/1000 | Detect FN with Western blot | Abcam, USA |
| β-tubulin | Mouse monoclonal<br>antibody-HRP conjugated | 1/5000 | Western blot internal reference | Affinity, China |
| β-actin | Rabbit monoclonal<br>antibody-HRP conjugated | 1/5000 | Western blot internal reference | ZSGB-BIO,<br>China |
| IgG(H+L) | Peroxidase-Conjugated<br>Goat anti-Mouse | 1/5000 | Label NL1 with Western blot | ZSGB-BIO,<br>China |
| IgG(H+L) | Peroxidase-Conjugated<br>Rabbit anti-Goat | 1/5000 | Label NL2 with Western blot | ZSGB-BIO,<br>China |
| IgG(H+L) | Peroxidase-Conjugated<br>Rabbit anti-Rabbit | 1/5000 | Label NL2 with Western blot | ZSGB-BIO,<br>China |

**TABLE 2:**
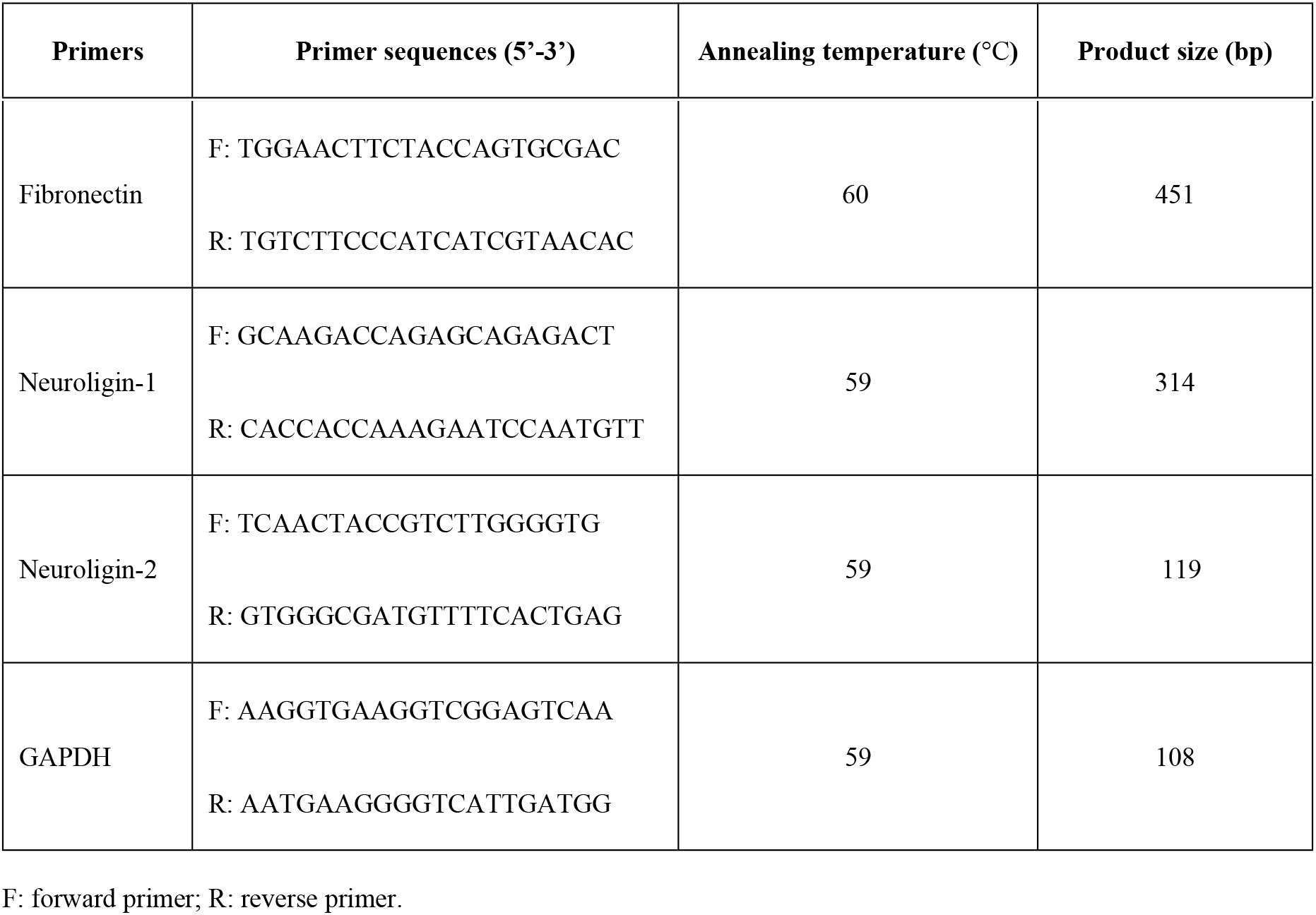
Detailed information of primers.

### Western blot analysis

Samples were lysed in ice-cold complete lysis buffer (RIPA: PMSF: protease inhibitor cocktail=100:1:1, Beyotime, Shanghai, China). The proteins were then quantified using the BCA Protein Assay Kit. Each plate of protein samples of the corresponding concentration was separated by an 8% SDS-PAGE gel and then transferred to a 0.45-μM polyvinylidene difluoride (PVDF) membrane. Membranes were first blocked with 5% nonfat milk in TBST for 2 hours at room temperature and then incubated with primary antibodies (Table 1) at 4°C overnight. After TBST washing, species-matched secondary antibodies (Table 1) were sequentially added for another 2 hours at room temperature. Finally, proteins were detected using an enhanced chemiluminescence (ECL) kit (Millipore, MA, USA) and visualized using the Chemidoc™ system (Bio-Rad). To verify the abundance of FN and NLs, ImageJ Image Processing Software was used to quantify the gray value of each sample blot, and then the relative gray values (FN, NL1, NL2/β-actin, β-tubulin) were obtained.

### Analysis of FN and NL gene expression by qRT-PCR

Total RNA was extracted using TRIzol Reagent (Invitrogen, USA) according to the manufacturer’s instructions to isolate total RNA from the colon samples and every plate of treated cells. The quantity of RNA was assessed spectrophotometrically by a NanoDrop OneC Microvolume UV-Vis Spectrophotometer, and the OD value at 260/280 for RNA ranged between 1.9 and 2.1. Reverse transcription by Tgradient 96 PCR was carried out using 500 ng of total RNA and PrimeScriptTM RT Master Mix kit according to the manufacturer’s protocol. The amplification of cDNA was performed according to the instructions of the SYBR Premix Ex TaqTM kit, and qRT-PCR was performed using an instrument with a Roche LightCycler 480 system. Within the instrument, the reaction mixture was first incubated at 95°C for 30 sec to denature the template DNA. Amplification was then performed for 40 cycles, each involving denaturation at 95°C for 30 sec, annealing for 30 sec at 59°C (NLs and GAPDH) or 60°C (FN), and elongation at 65°C for 15 sec. The relative gene expression data were analyzed by the 2^−Δ Δ Ct^ method.[17] The 0 μg/mL group served as a negative control, and 2^−Δ Δ Ct^ was calculated as the relative expression for further analysis. The expression of GAPDH in each sample was used as an internal control.

### Statistical analyses

All data shown in this study were analyzed with Graph Pad Prism® 5 software (La Jolla, CA, USA) and excel. One-way analysis of variance (ANOVA) with Tukey’s multiple comparison test was used for comparisons among three or four groups, and unpaired t-tests were used for comparisons between two groups. The original index data were standardized by Zero-mena normalization. All results are expressed as the mean values (±SD), and *P* values < 0.05 were considered statistically significant.

## Results

We have repeated our previous studies[13, 14] and confirmed that gene dysplasia of NLs and FN are associated with HSCR. HSCR is divided into three segments: aganglionic segments, transitional segments and normal tissue. NL1 and NL2 expression gradually increased in the three sections, while FN showed the opposite effect (data not shown). Previous studies suggest that there must be a relationship among the three, and the imbalance in their expression must be associated with HSCR.

### Downregulation of FN and the correlated expression of NLs in the enteric nervous system of embryonic rats

A WB assay was performed to investigate the expression of FN, NL1 and NL2 in the later embryonic stage with the aim of confirming the temporal increasing trend of NL1 and NL2 expression and the temporal decreasing trend of FN expression. Compared with the expression levels in E16 rats, the protein expression of NL1 and NL2 at E18, E20, and Ep0 were significantly increased, while the expression levels of FN were significantly decreased (P<0.05) (Fig 1A and B). And the results were consistent with those of the q-PCR analysis (P<0.05) (Fig 1C). Fig 1 reveals a temporal trend in which the expression of NL1 and NL2 gradually increased while FN gradually decreased during the development of ENS in the later embryonic stage, which meant that there was downregulation between FN and the correlated expression levels of NL1 and NL2 during the development of the embryonic ENS.

**Fig 1.**
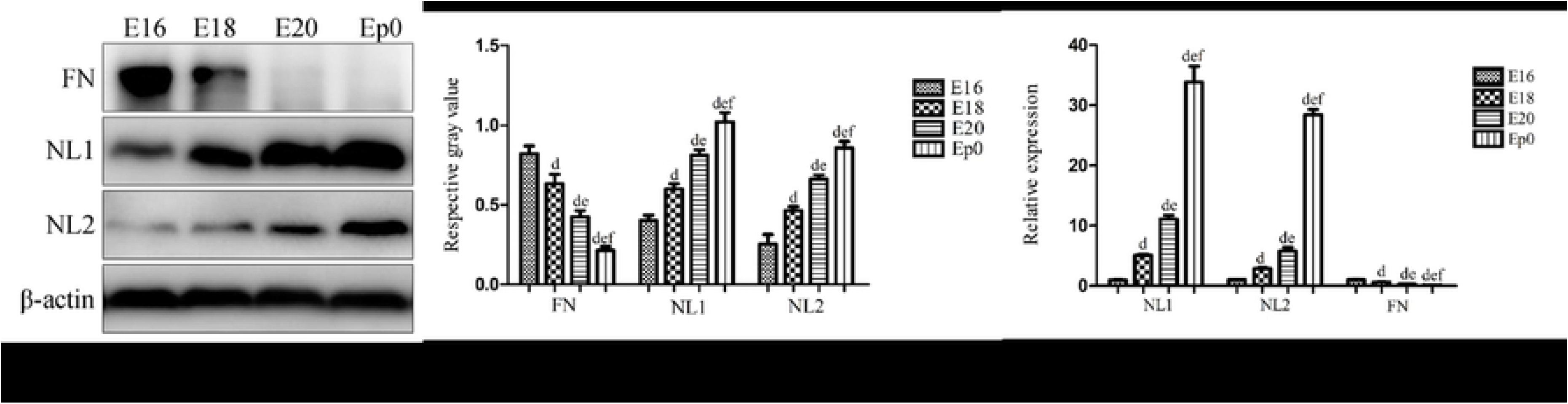
Western blot and qRT-PCR images showing that FN, NL1 and NL2 are present at time points E16, E18, E20, and Ep0. (A) Western blots of intestinal samples isolated from embryonic rats at E16, E18, E20 and Ep0 and then incubated with NL1, NL2 and FN antibodies. (B) Quantitative graphs of the results in A, showing a significant increase in NL1 and NL2 and a significant decrease in FN on different embryonic days (P< 0.05). (C) Relative expression of NL1, NL2 and FN mRNA on different embryonic days. Comparisons of the expression levels showed that NL1, NL2 and FN mRNA increased gradually along the time points of E16, E18, E20, and Ep0, and comparisons within groups were statistically significant (P< 0.05). ^d^P <0.05 versus E16, ^e^P < 0.05 versus E18, and ^f^P < 0.05 versus E20.

### Functional study of the downregulatory effects between FN and NLs Suppression of FN can promote NL expression

To confirm the direct relationship between FN and NLs, we transfected PC12 cells with NL1, NL2 and FN siRNA, referred to as knockdown (KD) lines of NL1-KD, NL2-KD and FN-KD, respectively. Then, the effect of genetic deletion of FN on NL1 and NL2 was detected by transfecting siRNA into PC12 cells. As shown in Fig 2C and 3C, F, the results of qRT-PCR demonstrated that the expression of FN, NL1 and NL2 were significantly decreased in the KD group compared with the negative control (NC) group, illustrating that the gene was successfully silenced with the siRNA technique (P<0.05). Moreover, the levels of NL1 and NL2 (Fig 2A and B) were significantly higher in the FN-KD group than that in the NC and control (Ctr) groups, while the FN was the opposite, indicating that FN inhibits the expression of NL1 and NL2 (P<0.05).

**Fig 2.**
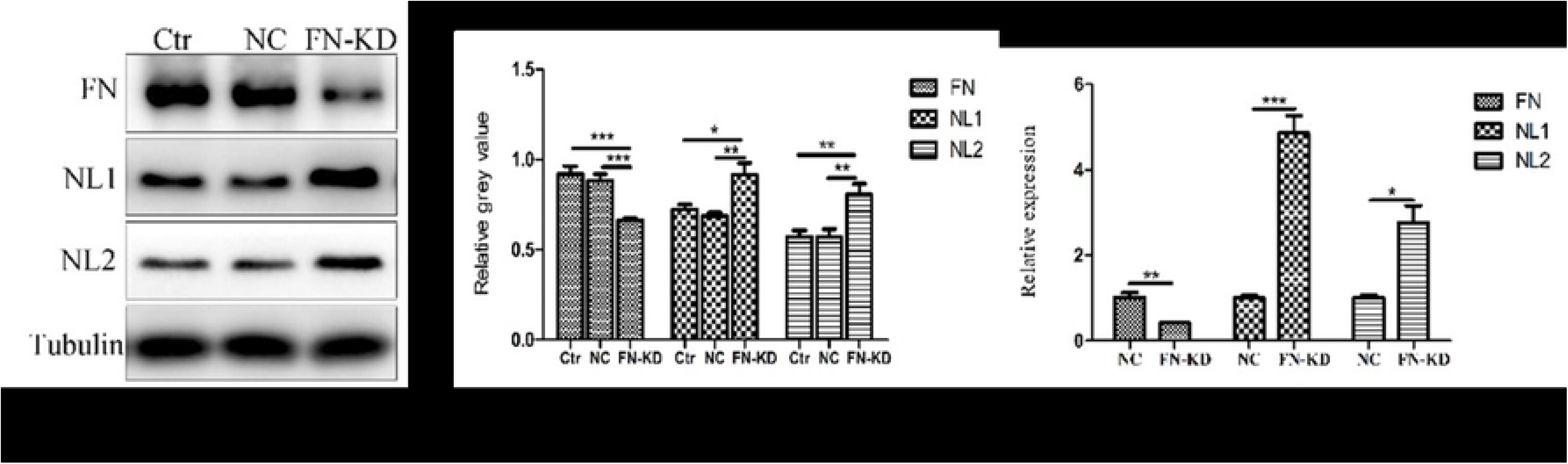
Downregulatory effects of FN on NLs studied in PC12 cell cultures in which the FN gene was knocked down with the siRNA technique. (A) Western blots of PC12 cell lysates in the FN-KD, NC and Ctr groups incubated with NL1, NL2 and FN antibodies, respectively. (B) Quantitative graphs of the results in A, showing a significant increase in NL1 and NL2 in the FN-KD group compared with the NC and Ctr groups. (C) Relative expression of NL1 and NL2 mRNA significantly increased in the FN-KD group compared with the NC group. *, P<0.05; **, P<0.01; ***, P<0.001.

**Fig 3.**
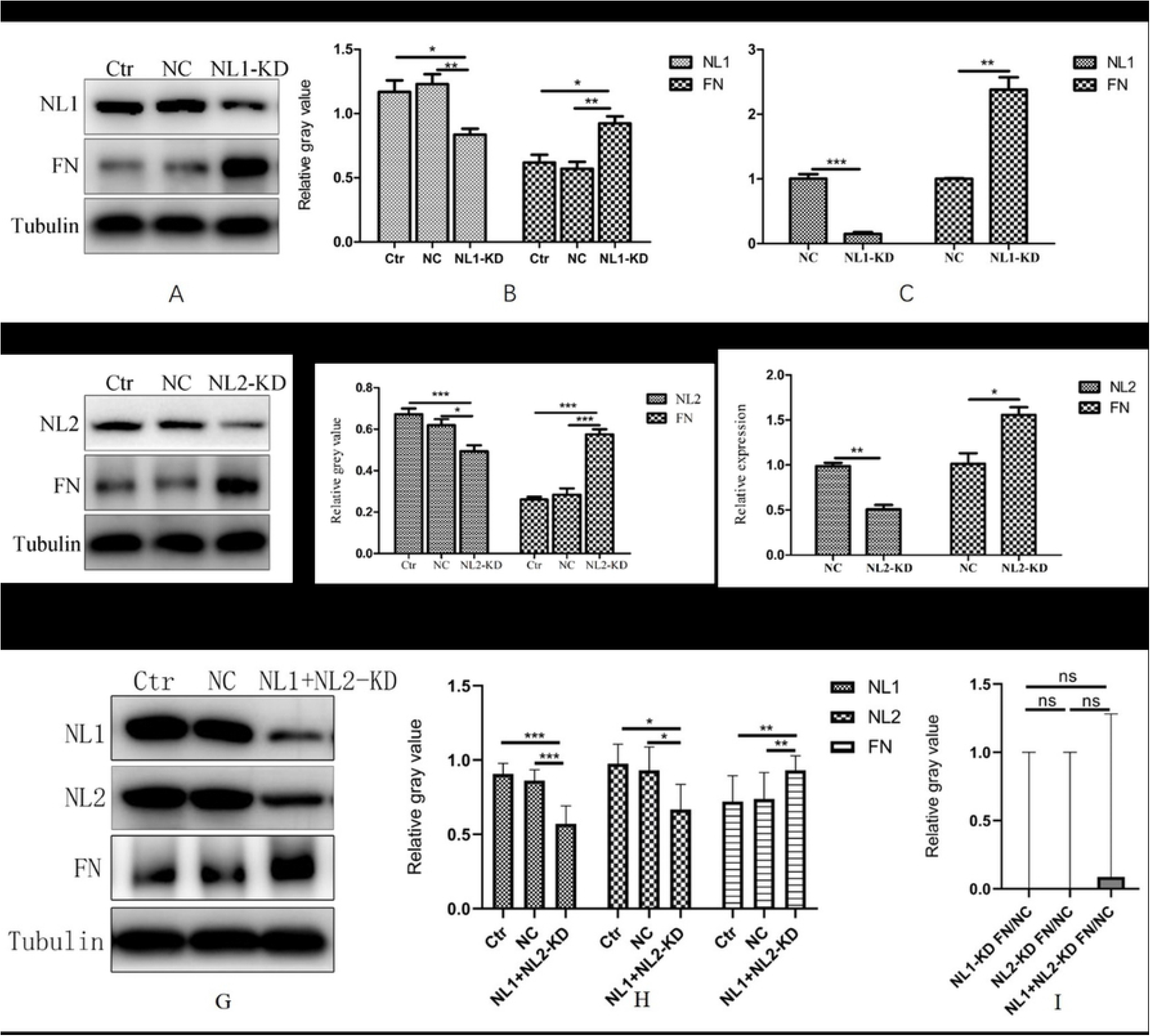
Downregulatory effects of NL1 and NL2 on FN in PC12 cell cultures in which the NL1 and NL2 gene was knocked down with the siRNA technique respectively. (A, D and G) Western blots of PC12 cell lysates in the NL-KD, NC and Ctr groups incubated with NL1, NL2 and FN antibodies, respectively. (B, E and H) Quantitative graphs of the results in A, D and G, showing significant changes in FN in the NL-KD group compared with the NC and Ctr groups. (C, F) Relative expression of FN mRNA was significantly changed in the NL-KD group compared with the NC group. (I) There was no significance on relative Z-score gray values of FN/NC protein among NL1-KD, NL2-KD and NL1+NL2-KD groups. *, P<0.05; **, P<0.01; ***, P<0.001; ns, not significant.

### Suppression of NL1 and NL2 can promote FN expression

As shown in Fig 3, WB analysis of the NL1 and NL2 siRNA-infected cells indicated that the expression of FN was much higher in the NLs-KD group than in the Ctr group and the NC group (P<0.05). The results of RT-PCR analysis were consistent with the WB results. After downregulating NLs, FN exhibited a reversed expression pattern, concluding that NL1 and NL2 can inversely affect the expression of FN.

With aim to evaluate the effect of the combination influence of NL1 and NL2 on the expression of FN, PC12 cells were simultaneously transfected with both NL1 and NL2 siRNA and their protein levels were evaluated. As shown in Fig 3, western blot analysis demonstrated that there was no significance on the expression of FN among NL1-KD, NL2-KD and NL1+NL2-KD groups, which means that there was no combination influence of NL1 and NL2 on the expression of FN.

### Negatively correlated relationship between FN and NLs in 293T cells

Our previous studies on HSCR have proved that FN expression in aganglionic segments, transitional segments and ganglionic segments showed a diminishing trend, while the expression of NL1 and NL2 was just the opposite.[18] Then, FN1 recombinant proteins (FN1rp), NL1 recombinant proteins (NL1rp) and NL2 recombinant proteins (NL2rp) were used to examine whether there was a linear relationship among FN, NL1 and NL2. High-level expression models of FN and NLs were produced by recombinant proteins, and their effects on the expression of NLs and FN were observed.

WB and RT-PCR were performed in 293T cells to measure the expression of FN and NLs for exploring the putative relationship between them. To investigate the optimal stimulation time, 20 μg/mL FN and 0.2 μM NLsrp were selected to stimulate for 0, 0.5, 1, or 4 hours. As depicted in Fig S1 and S2, the results demonstrate that the inhibitory effect of FN and NLs was the best when stimulated for 0.5 hours, which was selected as the basis for later experiments.

The relative mRNA expression levels along the four RGD groups (0, 5, 10, 20 μg/mL) indicate a temporal trend of gradually decreasing expression of NL1 and NL2 (Fig 4C). The abundance of transcripts encoding FN was significantly lower under each increasing concentration group, which means the relative expression of FN decreased as the recombinant protein concentrations increased (0 μM > 0.2 μM > 1 μM > 5 μM) in both NL1-FN and NL2-FN groups (Fig 4F).

**Fig 4.**
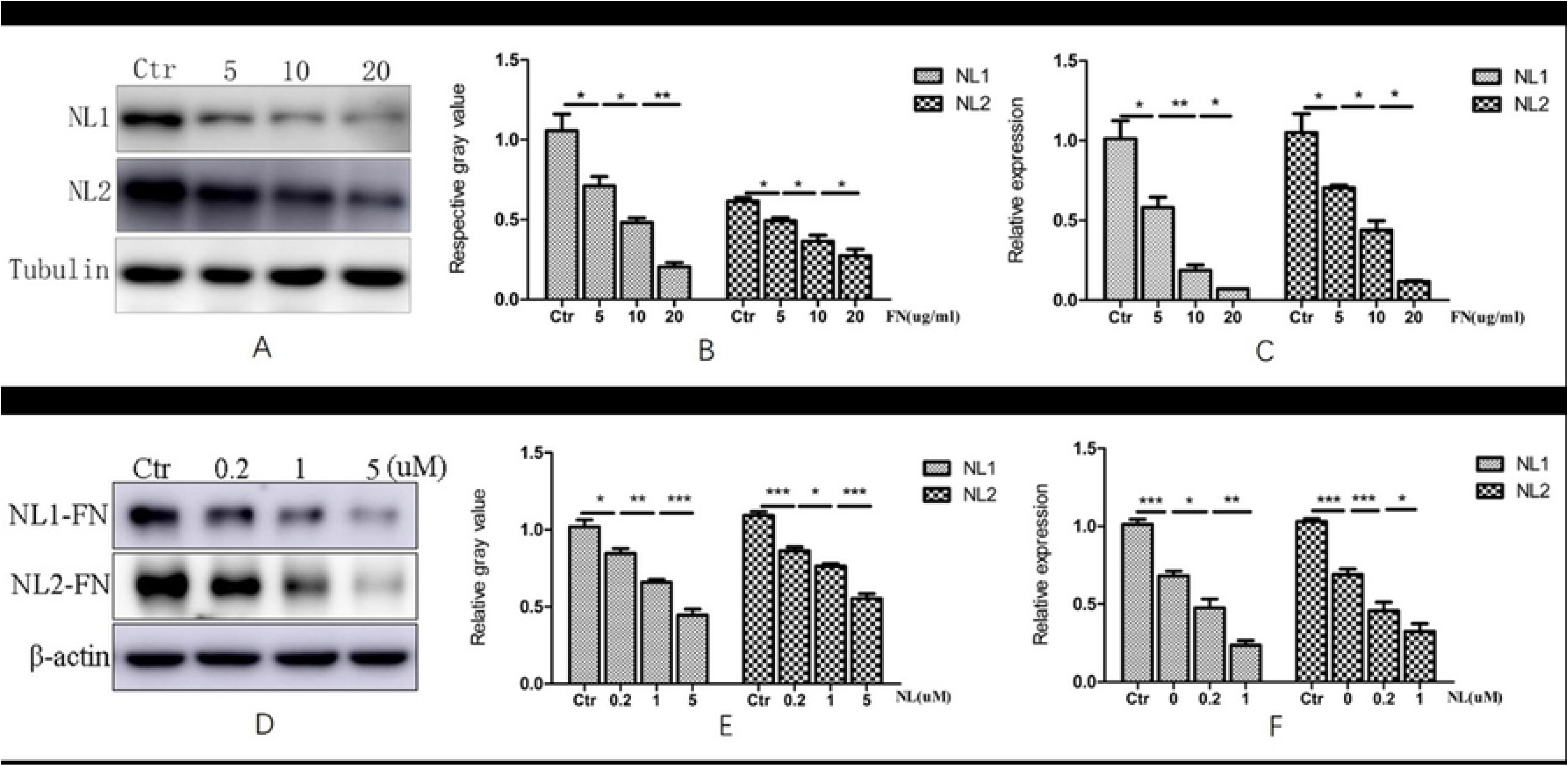
Western blot and qRT-PCR images showing that NL1, NL2 and FN are present with various concentrations of FN and NLs recombinant proteins. (A, D) Western blots of protein lysates from cultured 293T cells treated with various concentrations of FN and NLs recombinant proteins for 0.5 hours respectively. (B) Quantitative graph of the results in A and D, showing significantly decreased NL1, NL2 and FN with increased concentrations of FNrp and NLsrp. The differences in the relative gray values among the four groups (5 μg/mL vs Ctr; 10 μg/mL vs 5 μg/mL; 20 μg/mL vs 10 μg/mL) (0.2 μM vs. Ctr; 1 μM vs. 0.2 μM; 5 μM vs. 1 μM) were statistically significant (P<0.05). (C) Relative mRNA expression in 293T cells treated with various concentrations of recombinant proteins. The mRNA expression gradually decreased with increasing concentrations of FNrp and NLsp, and the comparisons within groups were significant(P<0.05). Ctr, control. *, P< 0.05. **, P< 0.01; ***, P<0.001.

Furthermore, WB analysis was performed to detect the protein levels of NL1, NL2 and FN in the four groups with increasing FN (0-20 μg/mL) and NLs (0-5 μM) concentrations. And the results of the immunoreactivity of proteins demonstrated by WB were consistent with the mRNA expression levels detected by RT-PCR (Fig 4A, B and D, E).

## Discussion

To date, 15 genes represented by RET have been implicated in HSCR development, only approximately 30% of which have mutations, suggesting the involvement of other genes or the abnormal ECM.[7, 19–22] Currently, it is widely accepted that the surrounding microenvironment is closely related to the corresponding nervous system, such as that in the intestinal tract. NL2 is similar in structure and sequence with NL1,[23] and it has been confirmed that NL1 and NL2 have similar activities *in vitro* in mice,[24] while NL1 is only expressed at excitatory synapses and NL2 at inhibitory synapses *in vivo*.[25] 293T cells, a human embryonic renal epithelial cell line, is widely used in cell biology for their reliable growth and propensity for transfection and the rat PC12 cell line was widely used in studies of neuronal disease, such as Alzheimer’s disease, Huntington’s disease and HSCR.[26, 27] In the current studies, PC12 cells and 293T cells were used to determine that high expression of FN can inhibit both NL1 and NL2, which means that the abnormal expression of FN can affect the expression of NLs on the postsynaptic membrane and that FN can directly affect the expression of genes as a member of the ECM environment. Given the subject of inducing or reducing the expression of NL, we found that NL1 and NL2 have similar activities *in vitro* in human and rat cells. Therefore, we speculated that there is a negative regulatory relationship between NLs and the ECM, and both NL1 and NL2 have similar effects on this regulation. Furthermore, we examined the expression of NLs (both NL1 and NL2) in 293T cells treated with four increasing concentrations of FN (0, 5, 10 and 20 μg/mL), which resulted in linearly decreasing expression of NL1 and NL2, and the result was consistent with that of the HSCR human model.[28] This experiment verified the negative correlation between the expression of FN and NL1/NL2 from the cellular machinery, and further studies relate FN, NL1 and NL2 with a negative linear correlation. The results provide an effective biological mechanism for future medical treatment. By transfection with siRNA, we confirmed that the mutual inhibition between FN and NLs was achieved by inhibiting their gene expression.

FN expression gradually increased from the ganglionic segment to the transitional segment and further to the aganglionic segments.[14] Similarly, increased accumulation of FN has been found in both the interfascicular and intrafascicular compartments in progressive bladder outlet obstruction.[29] We concluded that overexpression of FN plays a critical role in causing stenosis or obstructive disease. From this view, the lesions or immature intestinal environment of the HSCR intestinal microenvironment were simulated with increasing FN protein concentrations. The expression levels of NL1 and NL2 are low in HSCR, and their expression presented from highest to lowest in the ganglionic, transitional and aganglionic segments, while the expression patterns of FN were exactly opposite.[28] During embryonic growth and development, the ENS migrates in a rostral to caudal direction and finally settles in the entire fetal bowel to gradually develop and mature. In the process of distal migration and colonization of the ENS, we hypothesize that FN expression gradually decreases to prevent excessive migration, while NL expression gradually increases to mature the ENS. The decreased expression of FN further promotes the expression of NLs, that is, the development of the ENS, or the further expression of NLs inversely inhibits the expression of FN. Perhaps the balance between decreased FN expression and increased NLs plays an important role in ENS morphogenesis and functional maturity.

Moreover, there is indeed a certain regulatory relationship between the surrounding microenvironment and the ENS. In addition, our current studies have confirmed that there is a negative dose-dependent correlation between the expression of FN and NLs. Therefore, our further studies will focus on whether the *in vitro* induction of FN and NL expression can improve or cure the effects of HSCR in mouse models. We may be able to use the optimal physiological dose to achieve a cure or reduction in the symptoms of NIMs.

## Supporting information

**Fig S1.**
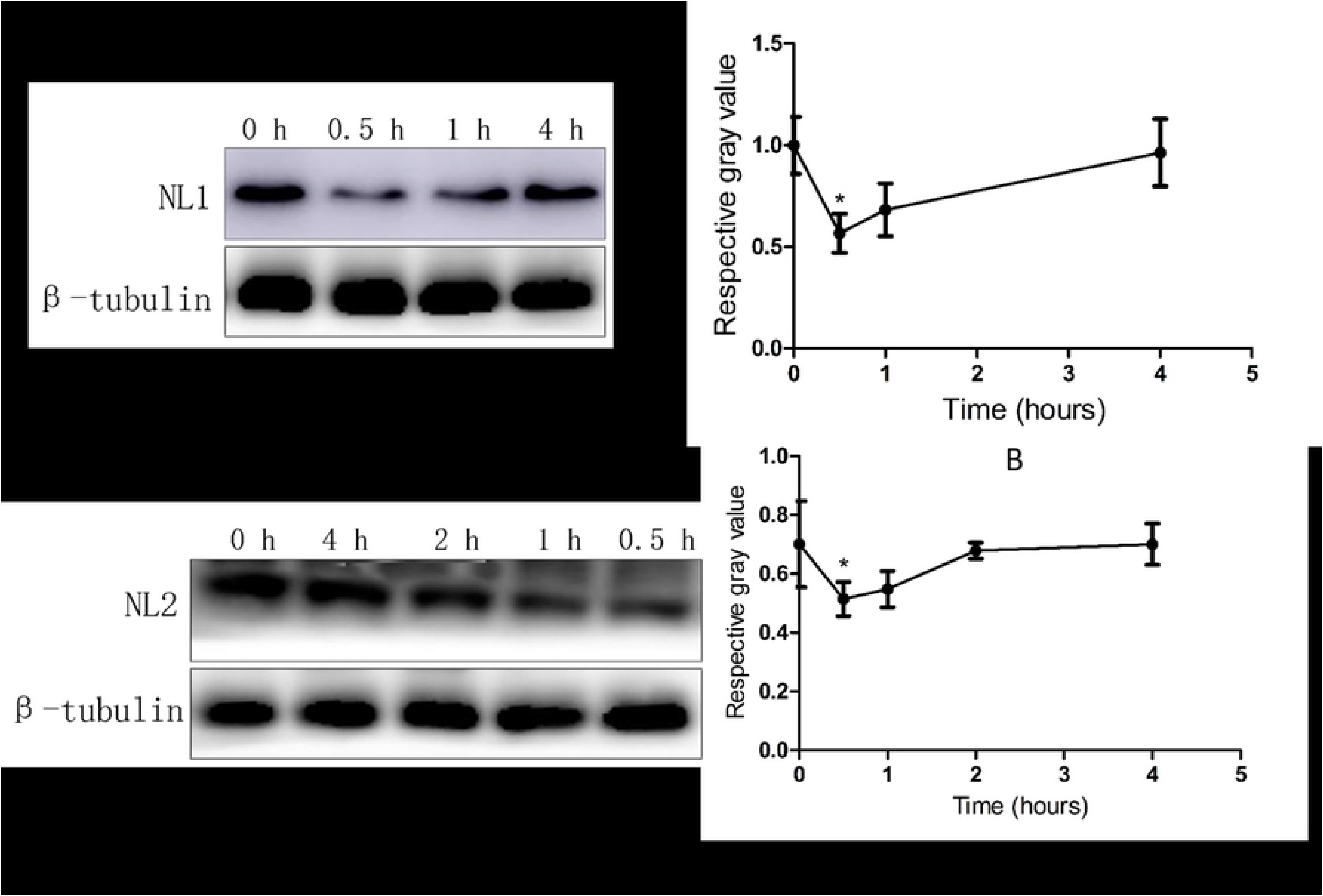
Western blots of NL1 and NL2 immunoreactivities of protein lysates from 293T cells treated with FN immediately or at various time points. (A) The levels of NL1 were dynamically changed. (B) Quantitative graphs of the results in A, showing the time course of NL1 levels during and after FN treatment. (C) The levels of NL2 were dynamically changed. (D) Quantitative graphs of the results in C, showing the time course of NL2 levels during and after FN treatment. *, P < 0.05.

**Fig S2.**
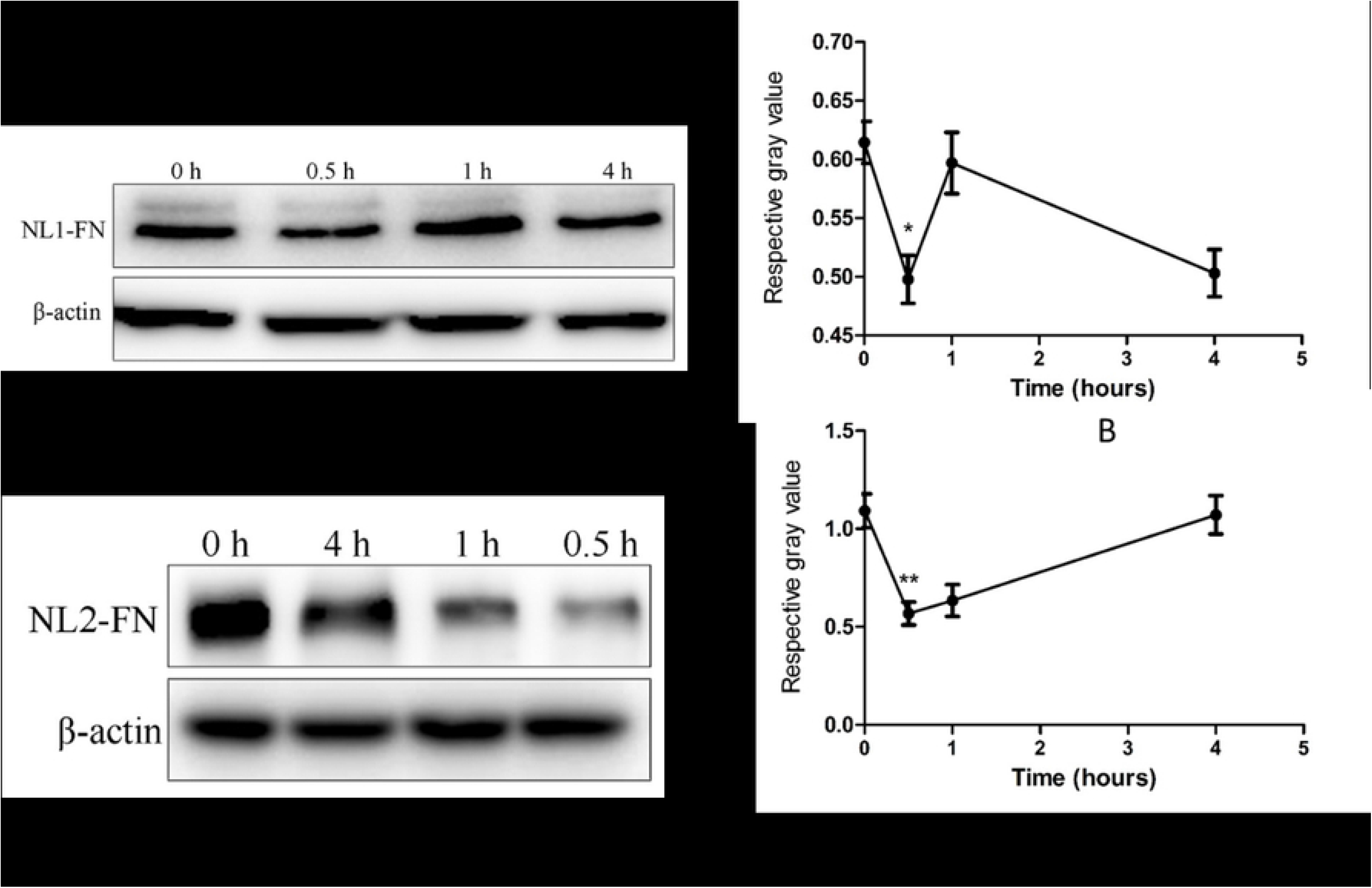
(A) Western blots of protein lysates from 293T cells immediately or at various time points after NL1rp treatment showing that the level of FN at different periods was dynamically changed. (B) Quantitative graphs of the results in A, showing the time course of FN levels during and after NL1 treatment. (C) Western blots of protein lysate from 293T cells immediately or at various time points after NL2 treatment showing that the level of FN was dynamically changed. (D) Quantitative graphs of the results in C, showing the time course of FN levels during and after NL2rp treatment. *, P < 0.05; **, P<0.01. NL1-FN, the expression of FN treated with NL1rp; NL2-FN, the expression of FN treated with NL2rp.

## Notes

## Acknowledgments

This study was supported by the National Natural Science Foundation of China (Projects nos. 81471487 and 81270720) and the Science Foundation of Qilu Hospital of Shandong University (Project no. 2015QLQN32).

## Authorship contributions

Ni Gao, Conceived project, Design and execution of most experiments, Data acquisition and analysis, Writing—original draft, review and editing; Peimin Hou, Data acquisition and analysis, Writing; Qiangye Zhang, Data acquisition and analysis; Weijing Mu, Data acquisition and analysis; Jian Wang, Writing—review and editing; Dongming Wang, Writing—review and editing; Aiwu Li, Conceived project, Resources, Supervision, Funding acquisition, Writing—review and editing.

Compliance with Ethical Standards

## Ethical Approval

Our study was approved by the ethics committee of Qilu Hospital, Shandong University. All procedures performed in studies involving animals were in accordance with the ethical standards of the institution or practice at which the studies were conducted.

